# Galactose-modified duocarmycin prodrugs as senolytics

**DOI:** 10.1101/746669

**Authors:** Ana Guerrero, Romain Guiho, Nicolás Herranz, Anthony Uren, Dominic J. Withers, Juan Pedro Martínez-Barbera, Lutz F. Tietze, Jesús Gil

## Abstract

Senescence is a stable growth arrest that impairs the replication of damaged, old or preneoplastic cells, therefore contributing to tissue homeostasis. Senescent cells accumulate during ageing and are associated with diseases, such as cancer, fibrosis and many age-related pathologies. Recent evidence suggests that the selective elimination of senescent cells can be effective on the treatment of many of these senescence-associated diseases. A universal characteristic of senescent cells is that they display elevated activity of the lysosomal β-galactosidase this has been exploited as a marker for senescence (senescence-associated β-galactosidase activity). Consequently, we hypothesised that galactose-modified cytotoxic prodrugs will be preferentially processed by senescent cells, resulting in their selective killing. Here, we show that different galactose-modified duocarmycin (GMD) derivatives preferentially kill senescent cells. GMD prodrugs induce selective apoptosis of senescent cells in a lysosomal β-galactosidase (GLB1)-dependent manner. GMD prodrugs can eliminate a broad range of senescent cells in culture, and treatment with a GMD prodrug enhances the elimination of bystander senescent cells that accumulate upon whole body irradiation or doxorubicin treatment of mice. Moreover, taking advantage of a mouse model of human adamantinomatous craniopharyngioma (ACP), we show that treatment with a GMD pro-drug result selectively reduced the number of β-catenin-positive preneoplastic senescent cells, what could have therapeutic implications. In summary, the above results show that galactose-modified duocarmycin prodrugs behave as senolytics, suggesting that they could be used to treat a wide range of senescence-related pathologies.

## INTRODUCTION

Cellular senescence is a stress response that prevents the replication of old, damaged or transformed cells (Herranz & Gil 2018). Senescence can be induced by replicative exhaustion and also by a range of insults that includes oncogenic activation, genotoxic stress or irradiation. The defining feature of senescence is a stable cell cycle arrest, but senescent cells also undergo multiple phenotypic changes including alterations in their morphology, metabolic state or chromatin arrangement (Salama *et al.* 2014). In particular, senescent cells secrete a combination of extracellular factors, the so-called senescence-associated secretory phenotype or SASP, which is a prominent mediator of the patho-physiological effects of senescence (Kuilman & Peeper 2009; Coppe *et al.* 2010).

Despite that the acute induction of senescence limits fibrosis and protects against cancer progression, the abnormal accumulation of senescent cells with age or in diseased tissues is considered detrimental (Munoz-Espin & Serrano 2014). Interestingly, recent evidences drawn from genetic models have shown that eliminating senescent cells increases lifespan, improves healthspan and benefits the outcomes of a wide range of diseases (Baker *et al.* 2011; Baker *et al.* 2016; Childs *et al.* 2016; Childs *et al.* 2017). These studies have led to a collective effort to identify ‘senolytics’, drugs that selectively kill senescent cells. Several senolytics have been identified including dasatinib and quercetin (Zhu *et al.* 2015), piperlongumine (Wang *et al.* 2016), FOXO4 interfering peptides (Baar *et al.* 2017), HSP90 inhibitors (Fuhrmann-Stroissnigg *et al.* 2017) or the Bcl2 family inhibitors ABT-263 (navitoclax) and ABT-737 (Chen *et al.* 2015; Yosef *et al.* 2016; Zhu *et al.* 2016). Currently, Bcl2 family inhibitors have become the gold-standard on senolysis. Bcl2 family inhibitors eliminate a range of senescent cells *in vivo* and reproduce the effects observed in transgenic mice modelling senescence ablation (Ovadya & Krizhanovsky 2018). However, ABT-263, causes severe thrombocytopenia and neutropenia, what might complicate its use on the clinic. Moreover, it is becoming evident than different senolytics might be necessary to eliminate different types of senescent cells. Therefore, there is a need to identify additional drugs with senolytic properties.

An alternative strategy for targeting senescence, is to exploit properties that differentiate senescent from normal cells. In this regard, the senescence-associated β-galactosidase activity (SA-β-gal) is one of the more conserved and defining characteristics of senescent cells. Senescent cells present an increased lysosomal mass (Kurz *et al.* 2000). As a result, senescent cells display elevated levels of lysosomal enzymes such as β-galactosidase (encoded by GLB1, (Dimri *et al.* 1995)) or α-fucosidases (Hildebrand *et al.* 2013). Indeed, it has been shown that galacto-oligosacharide encapsulated nanoparticles (GalNP) preferentially release their content on senescent cells (Agostini *et al.* 2012). Consequently, this GalNP can be used in combination with different cargos to either image or kill senescent cells (Munoz-Espin *et al.* 2018).

Galactose-modification has been frequently used to improve the pharmacokinetic properties or the delivery of existing drugs. In addition, galactose modification can be used to generate pro-drugs that rely on *E. coli* β-galactosidase for controlled activation (Melisi *et al.* 2011). When combined with antibody-linked β-galactosidase, this approach is known as antibody-directed enzyme prodrug therapy (ADEPT), (Bagshawe 2006; Tietze & Schmuck 2011). In ADEPT, a conjugate of a tumour-specific antibody and an enzyme, such as β-galactosidase, is combined with the application of a hardly cytotoxic prodrug. By means of the enzyme in the conjugate, the prodrug is selectively cleaved in cancer cells leading to the formation of a highly cytotoxic compound. Several of these galactose-modified cytotoxic prodrugs have been described (Leenders *et al.* 1999). A class of such prodrugs are galactose-modified duocarmycin (GMD) derivatives (Tietze *et al.* 2006). Duocarmycins are a group of antineoplastic agents with low picomolar potency. They are thought to act by binding and alkylating double stranded DNA in AT-rich regions of the minor groove, (Boger *et al.* 1994; Tietze *et al.* 2006; Tietze *et al.* 2009), but alternative mechanisms of action have been proposed to account for the cytotoxic effects of duocarmycin dimers (Wirth *et al.* 2012).

Here, we investigated whether galactose-modified prodrugs can preferentially kill senescent cells. We have assessed the senolytic potential of several GMD derivatives and confirmed their senolytic potential in cell culture, *ex vivo* and *in vivo*. Given the increasing list of senescence-associated diseases and the positive effects associated with senolytic treatment, we propose GMD derivatives and more generally galactose-modified prodrugs are a new class of senolytic compounds with wide therapeutic promise.

## RESULTS

### A galactose-modified duocarmycin prodrug with senolytic properties

The natural antibiotic duocarmycin is a highly cytostatic compound (Boger & Johnson 1995). A series of glycosidic derivatives of duocarmycin have been previously developed to be used as prodrugs in the context of antibody-directed enzyme prodrug therapy (ADEPT) (Tietze *et al.* 2009; Tietze *et al.* 2010). Given that senescent cells display elevated levels of SA-β-Galactosidase activity, we hypothesize that galactose-modified cytotoxic prodrugs will be preferentially processed by senescent cells, resulting in their selective killing. To test this hypothesis, we took advantage of a galactose-modified duocarmycin (GMD) prodrug (referred as prodrug A) previously described (Tietze *et al.* 2009). We analysed the effects that a seco-duocarmycin analogue dimer (duocarmycin SA) and its galactose derivative (prodrug A) had on the survival of IMR90 ER:RAS cells, a model of oncogene-induced senescence (OIS). Activation of the ER:RAS fusion with 4-hydroxy-tamoxifen (4OHT) induces senescence in IMR90 ER:RAS cells (Georgilis *et al.* 2018). Treatment with duocarmycin SA was equally effective in killing normal and senescent cells, with the exception of a small selectivity towards senescent cells at the lower concentrations (Fig 1a). In contrast, when we treated IMR90 ER:RAS cells with prodrug A (differing only in the addition of two galactose groups that inactivate it), we observed the preferential elimination of senescent cells (Fig 1b). Duocarmycins are known to bind and alkylate DNA in AT-rich regions of the minor groove, and induce cell death in a way dependent of DNA replication (Boger *et al.* 1994; Tietze *et al.* 2006; Tietze *et al.* 2009) We checked that senescent cells were growth arrested at the time of the drug treatment (Sup Fig 1). This shows that the effect observed is not due to hyperreplication of cells during early stages of OIS and suggest that the prodrug might act by some of the alternative cytotoxic mechanisms described for duocarmycin dimers (Wirth *et al.* 2012). Treatment with prodrug A induced caspase 3/7 activity on senescent cells (Fig 1c), and the selective death of this cells was prevented with a pan-caspase inhibitor (Fig 1d). The above results suggest that GMD prodrugs can behave as senolytics by selectively inducing apoptosis on senescent cells.

**Figure 1.**
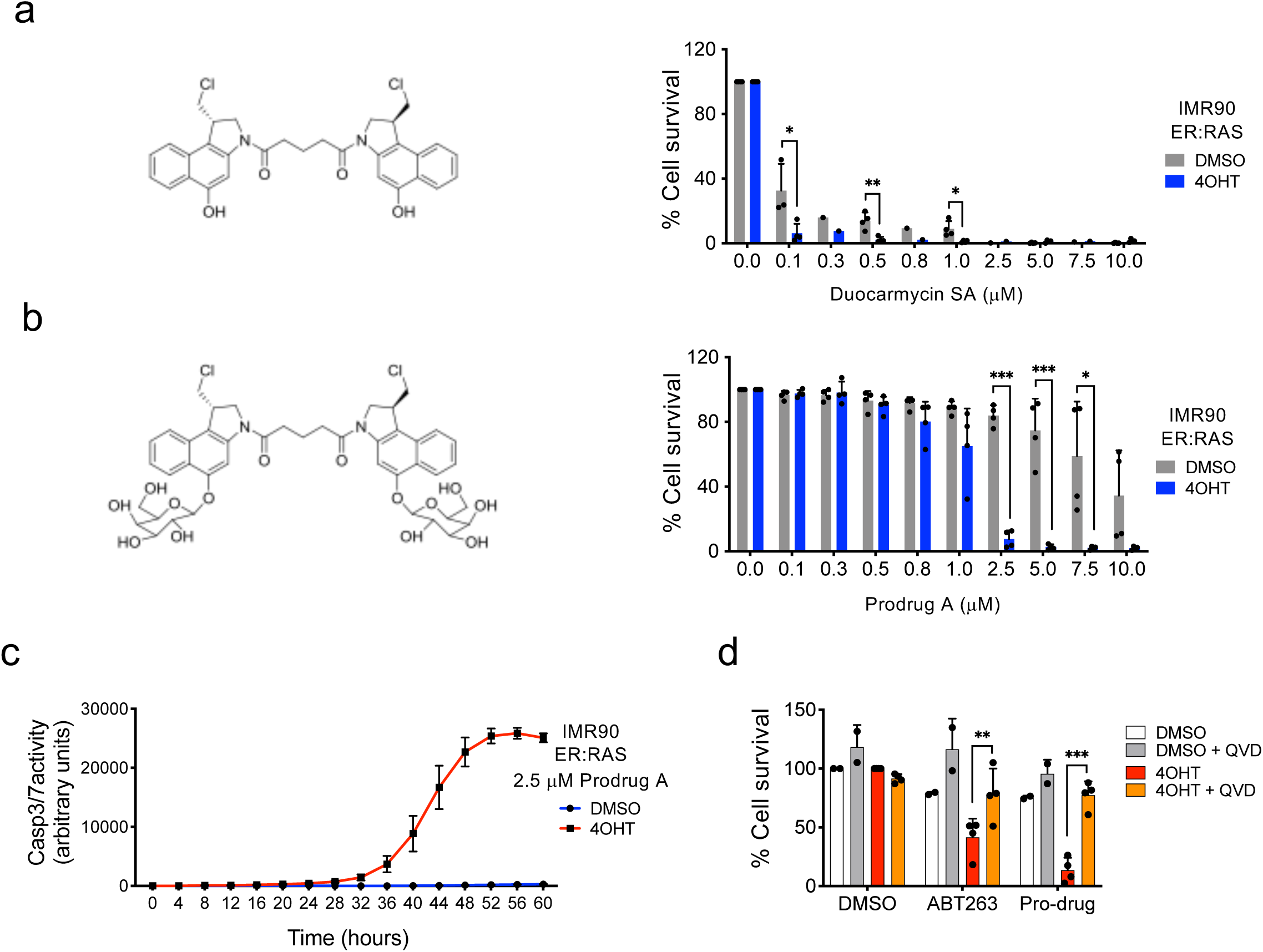
A galactose-modified duocarmycin prodrug with senolytic properties. **(a)** Molecular structure of seco-duocarmycin analog dimer (left). IMR90 ER:RAS cells were treated with DMSO or with 4OHT (4-hydroxytamoxifen) for 6 days to induce OIS. Cells were treated with the indicated concentrations of seco-duocarmycin analog dimer for 72 hrs. Cell numbers were quantified using DAPI staining and percentage of survival cells are plotted (right) (*n* = 4). **(b)** Molecular structure of a galactose-modified prodrug derivative of seco-duocarmycin analog dimer (referred as prodrug, left). Cell were treated with the prodrug for 72 hrs as described before (*n* = 4). Statistical significance was calculated using unpaired two-tailed Student’s *t*-tests. **(c)** Treatment of senescent cells with a GMD prodrug triggers caspase-3/7 activity. IMR90 ER:RAS were treated with 4OHT or vehicle (DMSO) for 6 days to induce senescence. 2.5 μM prodrug was then added together with NucLight Rapid Red reagent for cell labelling and Caspase-3/7 reagent for apoptosis (IncuCyte). Caspase 3/7 activity was measured at 4h intervals. **(d)** After 6-day treatment with 4OHT or vehicle (DMSO), IMR90 ER:RAS were treated with 1 μM ABT-263 or 2.5 μM Pro-Drug A for 72 hours in the presence or absence of the pan-caspase inhibitor QVD-OPh (*n* = 4). Statistical significance was calculated using two-way ANOVA (Tukey’s test). All error bars represent mean ± s.d; *n* represents independent experiments.; ns, not significant; \**P* < 0.05; \*\**P* < 0.01; \*\*\**P* < 0.001.

### Galactose-modified duocarmycin prodrugs are broad-spectrum senolytics

To understand the extent to which GMD prodrugs behave as senolytics, we assessed the effect that prodrug A has on several types of senescent cells. To this end, we took advantage of IMR90 cells and induced senescence by etoposide or doxorubicin treatment, irradiation, or serial passage. In all those instances, treatment with prodrug A resulted in the selective elimination of senescent cells (Fig 2a-d). Moreover, to evaluate whether the senolytic effects of prodrug A were restricted to IMR90 cells or also observed in other cell types, we took advantage of human mammary epithelial cells able to undergo OIS upon Ras activation (HMEC ER:RAS). Prodrug A was also able to selectively kill HMEC senescent cells (Fig 2e), suggesting that its senolytic effects were not cell type restricted. Finally, we wanted to examine whether the senolytic properties were specific of prodrug A, or the general concept (conversion of other cytotoxic drugs in galactose-modified prodrugs) was wider. To this end we took advantage of two previously described GMD prodrugs, JHB76B and JHB35B, (Tietze *et al.* 2009; Tietze *et al.* 2010). Both drugs were also effective in selectively eliminating senescent cells (Fig 2f and Sup Fig 2), suggesting that generation of galactose-modified prodrugs might be a general route to design senolytic compounds.

**Figure 2.**
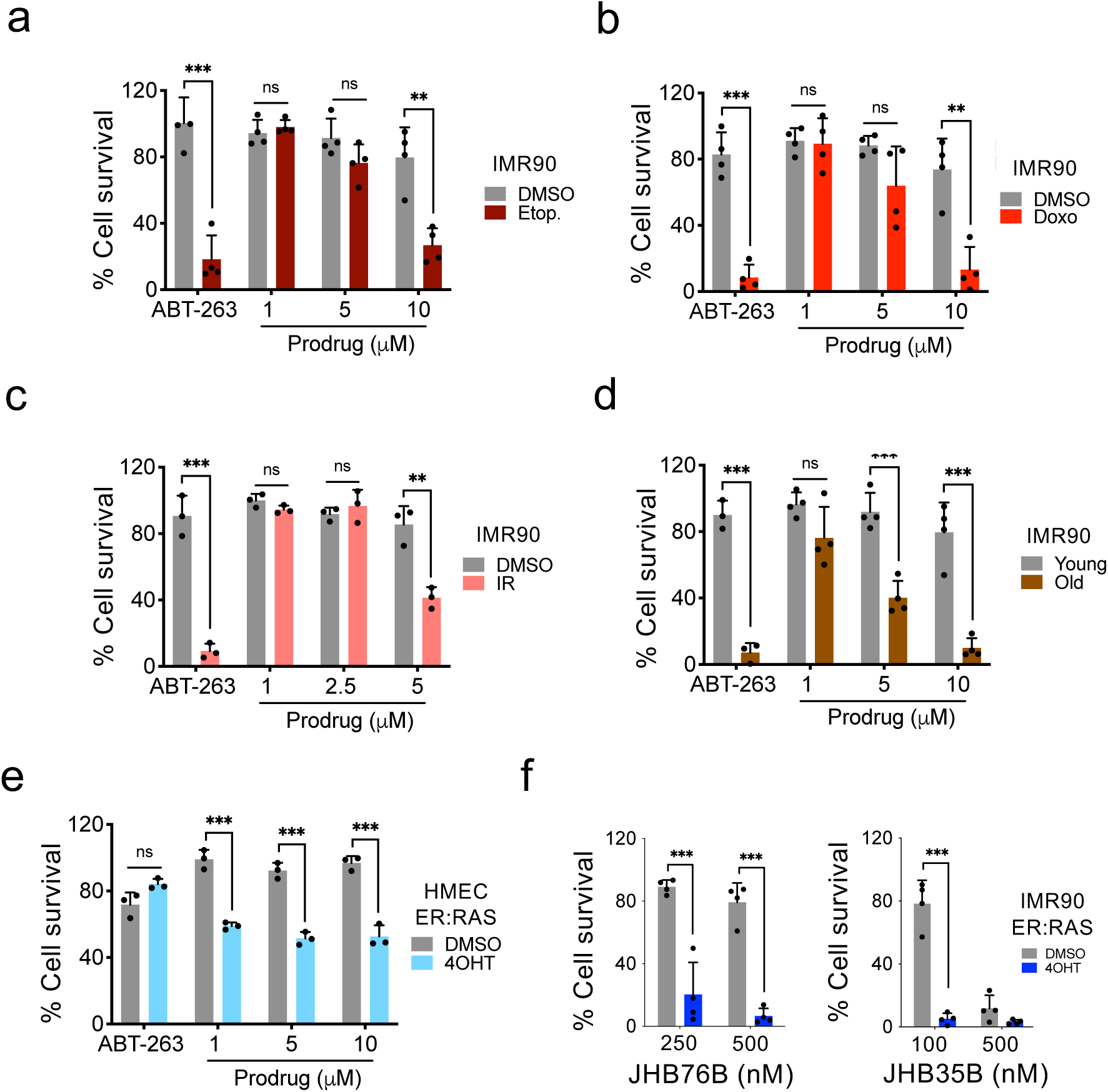
Galactose-modified duocarmycin prodrugs are broad-spectrum senolytics. **(a-d)** Quantification of cell survival after treatment with prodrug A in IMR90 undergoing different types of senescence. Senescence was induced by treatment with 50 μM etoposide (**a**, *n*=4), 0.5 μM doxorubicin (**b**, *n*=4) or 7.5 Gy irradiation (**c**, *n*=3). In (**d**) the effect of prodrug A on replicative senescence of IMR90 cells (passage 12 *vs* passage 22) (*n* = 4) was assessed. **(e)** Quantification of cell survival after treatment with prodrug A in HMEC ER:RAS, human mammary epithelial cells expressing hTERT that undergo senescence upon activation of ER:RAS by 4OHT treatment (*n* = 3). **(f)** Quantification of cell survival after treatment with two other galactose-modified duocarmycin derivatives, JHB76B and JHB35B, in the context of oncogene-induced senescence in IMR90 ER:RAS (*n* = 4). An extended version of this figure including additional drug concentrations is shown in Supplementary Figure 2. All statistical significances were calculated using unpaired two-tailed Student’s *t*-tests. All error bars represent mean ± s.d; *n* represents independent experiments.; ns, not significant; \**P* < 0.05; \*\**P* < 0.01; \*\*\**P* < 0.001.

### Senolytic properties of prodrug A depend on the lysosomal β-galactosidase

We had initially hypothesized that GMD prodrugs could behave as a senolytic due to the higher SA-β-Galactosidase activity of senescent cells. Increased β-Galactosidase on senescent cells is due to an increase in lysosomal mass (Kurz *et al.* 2000) resulting in higher activity of the lysosomal β-Galactosidase (encoded by GLB1) (Lee *et al.* 2006). To understand if this was the case and senolytic activity of GMD prodrugs is dependent on SA-β-Galactosidase, we took advantage of three independent shRNAs to knockdown *GLB1*. Knockdown of *GLB1* in IMR90 ER:RAS cells, resulted in decreased SA-β-Galactosidase activity, but it did not impact the growth arrest or the induction of p16^INK4a^ observed during OIS (Fig 3a-d). Taking advantage of these cells, we observed that *GLB1* knock down did not affect the senolytic potential of ABT-263 but ablated the ability of prodrug A to selectively kill senescent cells (Fig 3e). In summary, our data suggests that GMD prodrugs trigger apoptosis of senescent cells in a GLB1-dependent manner.

**Figure 3.**
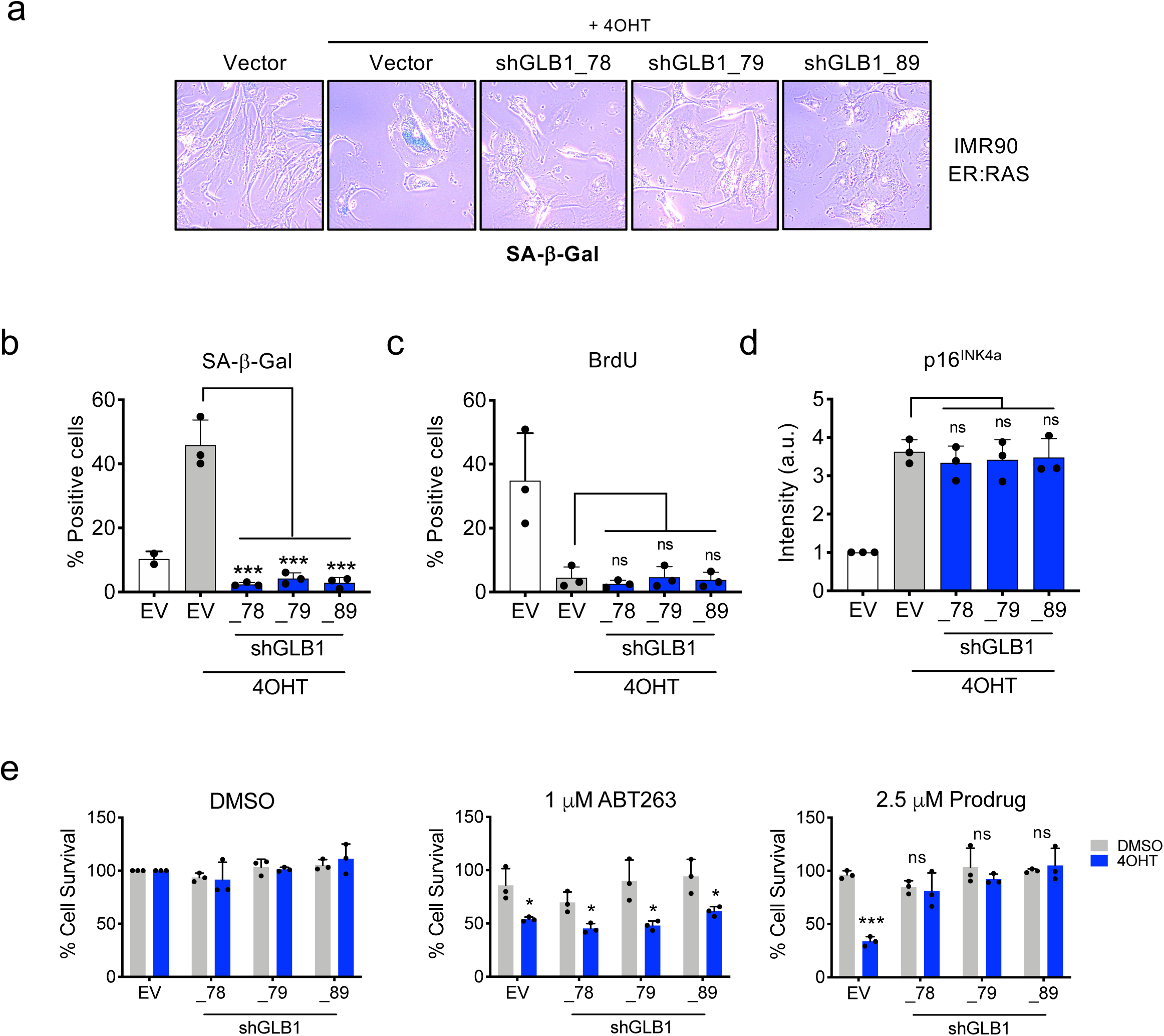
The senolytic properties of prodrug A depend on the lysosomal β-galactosidase (GLB1). **(a)** Representative pictures of cytochemical SA-β-Gal staining in IMR90 ER:RAS infected with different shRNAs against *GLB1* or an empty vector and **(b)** quantification (*n* = 3). Statistical significance was calculated using one-way ANOVA (Dunnett’s test). **(c-d)** IMR90 ER:RAS cells with reduced β-galactosidase expression undergo senescence as assessed by BrdU incorporation (**c**) and p16^INK4a^ staining (**d**). *n* = 3. Statistical significance was calculated using one-way ANOVA (Dunnett’s test). (**e**) Quantification of cell survival of senescent and control IMR90 ER:RAS infected with different shRNAs targeting *GLB1* or an empty vector and treated with ABT-263, prodrug A or vehicle (DMSO) for 3 days (*n* = 3). Statistical significance was calculated using two-tailed, Student’s *t*-test. All error bars represent mean ± s.d; *n* represents independent experiments; ns, not significant; \**P* < 0.05; \*\**P* < 0.01; \*\*\**P* < 0.001.

### Prodrug A eliminates bystander senescent cells *in vivo*

Chemotherapy and radiotherapy are amongst the most common anti-cancer treatments. Irradiation, chemotherapy and even some targeted anti-cancer drugs, all induce senescence (Schmitt *et al.* 2002; Wang *et al.* 2017). Although induction of tumour senescence explains the anti-cancer properties of these treatments, the generation of bystander senescent cells is responsible for their side effects (Demaria *et al.* 2016). To assess whether pro-drug A could eliminate these bystander senescent cells, we first irradiated mice and upon a latency period to allow for the accumulation of senescent cells, treated them with prodrug A, ABT263 or vehicle (Fig 4a). Treatment with prodrug A or ABT-263 resulted in a reduced presence of senescent cells in lung as assessed using SA-β-Galactosidase activity (Fig 4b).

**Figure 4.**
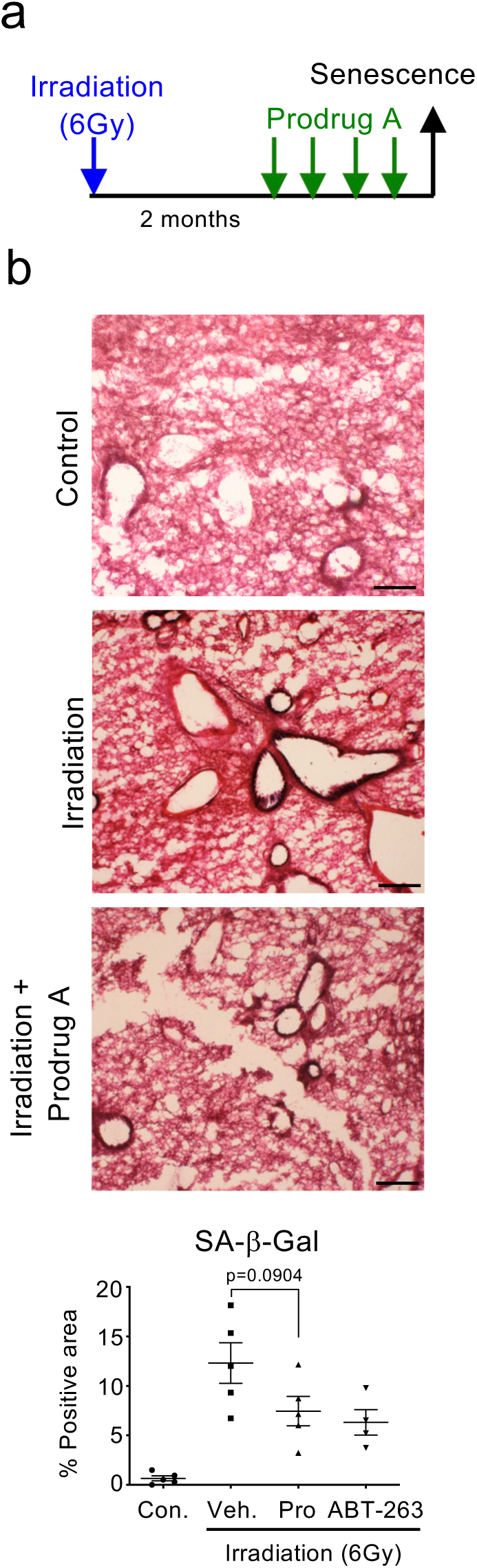
Galactose-modified duocarmycin prodrug eliminates bystander senescent cells accumulating *in vivo*. Prodrug A eliminates senescent cells accumulating after whole body irradiation. **(a)** Experimental design of the whole-body irradiation-induced senescence experiment. Mice (*n* = 4/5 per group) were irradiated with 6 Gray to induce senescence. 2 months later mice were treated with vehicle, Prodrug A (JHB75B) or ABT-263 for 4 consecutive days, before being culled for analysis. **((b)** Representative pictures of lung cryosections (top) and quantification of the lung area positive for SA-β-Gal staining (bottom). Statistical significance was calculated using unpaired Student’s *t*-test. Data represent mean ± s.d; *n* represents number of mice; ns, not significant; \**P* < 0.05; \*\**P* < 0.01; \*\*\**P* < 0.001.

### Galactose-modified prodrugs eliminate preneoplastic senescent cells

OIS is primarily considered as a tumour suppressive mechanism (Collado *et al.* 2005) but senescent cells present in the tumour microenvironment can also drive tumour progression (Gonzalez-Meljem *et al.* 2018). We have previously demonstrated in mouse models of adamantinomatous craniopharyngioma (ACP), a pituitary paediatric tumour, that clusters of cells that accumulate nucleo-cytoplasmic β-catenin are senescent and drive tumour progression in a paracrine manner (Gonzalez-Meljem *et al.* 2017). To understand if GMD prodrugs could eliminate these pro-tumourigenic senescent clusters, we used the *Hesx1*^*Cre/+*^;*Ctnnb1*^*lox(ex3)/+*^ ACP mouse model. Tumoural, cluster-containing embryonic pituitaries were cultured *ex vivo* with vehicle or prodrug A (Fig 5a). Treatment with prodrug A preferentially eliminated the β-catenin-accumulating senescent cell clusters, without affecting other cell types in the pituitary such as synaptophysin + cells (Fig 5b-d). Co-staining with an antibody recognizing cleaved caspase 3 showed that prodrug A predominantly induced apoptosis of senescent cluster cells (Fig 5e and Sup Fig 3). The above results suggest that GMD prodrugs can be also used to eliminate preneoplastic senescent cells.

**Figure 5.**
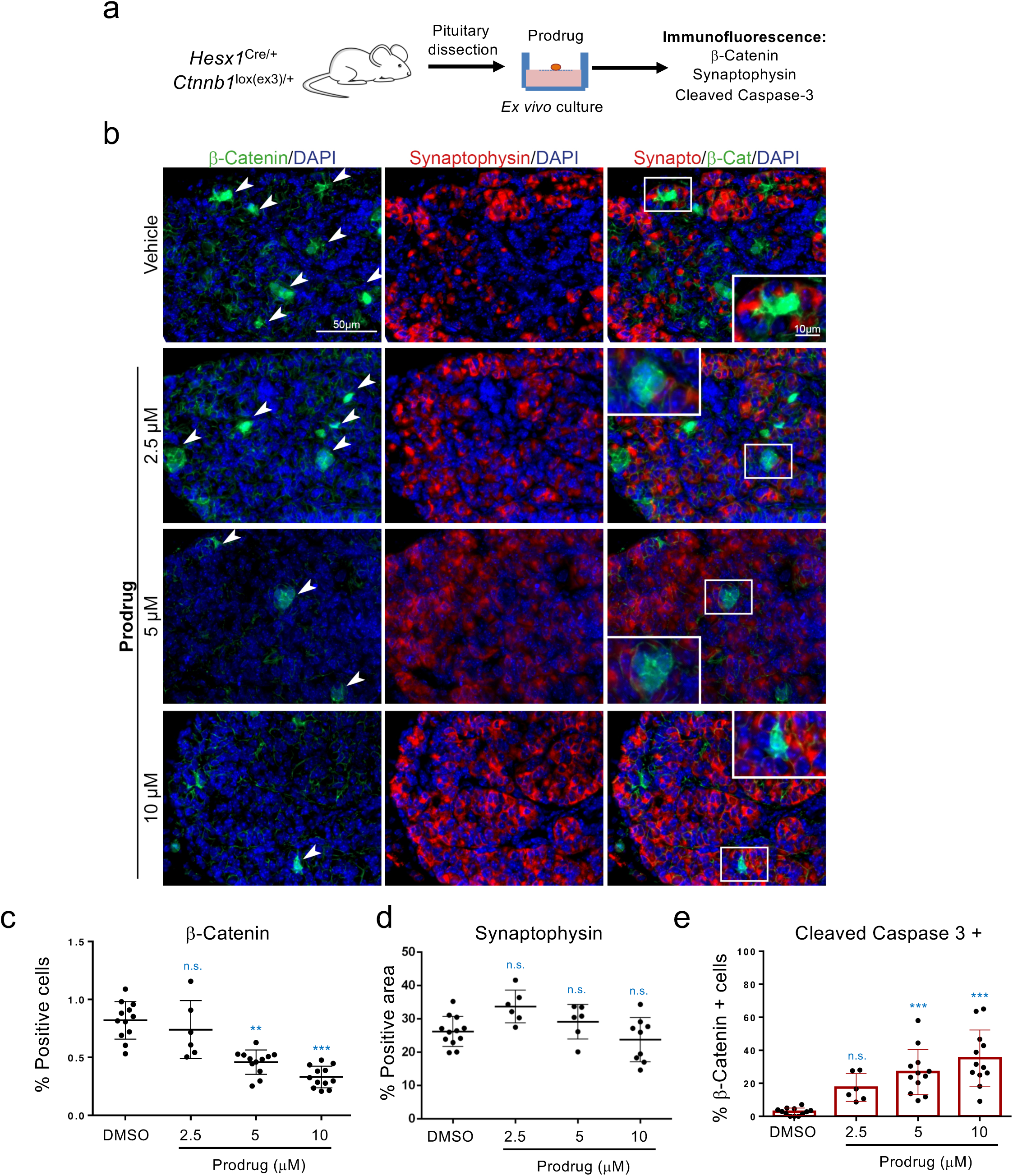
Galactose-modified duocarmycin prodrug eliminates preneoplastic senescent lesions. **(a)** Experimental design for the senolytic experiment in the *Hesx1*^*Cre/+*^;*Ctnnb1*^*lox(ex3)/+*^ mouse model of adamantinomatous craniopharyngioma (ACP). Tumoural pituitaries from 18.5dpc *Hesx1*^*Cre/+*^;*Ctnnb1*^*lox(ex3)/+*^ mice were cultured in the presence of Prodrug A at the indicated concentrations or vehicle (DMSO) and processed for analysis after 72 hours. **(b)** Immunofluorescence staining against β-catenin (green) and synaptophysin (red) is shown. Synaptophysin is a marker of the normal hormone-producing cells in the pituitary gland. Scale bar, 50μm. **(c)** Quantification of β-catenin-accumulating cells after treatment with different concentrations of pro-drug A or vehicle (*n* = 6-12). Statistical significance was calculated using non-parametric ANOVA with Dunn’s *post hoc* comparison. **(d)** Quantification of synaptophysin-positive cells after treatment with different concentrations of pro-drug A or vehicle (*n* = 6-12). Statistical significance was calculated using non-parametric ANOVA with Dunn’s *post hoc* comparison. (**e**) Quantification of β-catenin-accumulating cells positive for cleaved-caspase 3 after treatment with different concentrations of prodrug A or vehicle (*n* = 6-12). Statistical significance was calculated using non-parametric ANOVA with Dunn’s *post hoc* comparison. All error bars represent mean ± s.d; *n* represents number of pituitaries; ns, not significant; \**P* < 0.05; \*\**P* < 0.01; \*\*\**P* < 0.001.

## DISCUSSION

Recently, the use of genetic mouse models in which senescent cells can be ablated, has served to unveil important roles for senescence in health, disease and aging (Baker *et al.* 2011; Demaria *et al.* 2014; Baker *et al.* 2016). Consequently, drugs have been identified that are able to phenocopy the effects of selectively eliminating senescent cells. Several of these so-called senolytic drugs have been discovered, with Bcl2 family inhibitors such as ABT-263 (navitoclax) and ABT-737 (Chen *et al.* 2015; Yosef *et al.* 2016; Zhu *et al.* 2016) being the prototypical examples.

Here, we add galactose-modified duocarmycin (GMD) prodrugs as a new class of senolytic agents. These GMD prodrugs are converted to their corresponding duocarmycin drugs in a manner dependent on processing by β-galactosidase. Since senescent cells display elevated levels of lysosomal β-galactosidase (encoded by *GLB1*), GMD selectively affect senescent cells. In this manuscript, we present evidence showing that GMD prodrugs can eliminate multiple types of senescent cells, what is consistent with SA-β-galactosidase being a universal marker of senescence. Moreover, we show that GMD prodrugs are also capable of eliminating bystander senescent cells caused by anti-cancer therapies and preneoplastic senescent cells in mouse models. Given the promise that senolytics present for the treatment of age-related disease, and their associated benefits over healthspan and lifespan, we believe that this study provides the basis to specifically assess the potential benefots of GMD on ageing.

Previously, the potential to harness the elevated β-galactosidase activity of senescent cells have been exploited with galacto-oligosaccharide encapsulated nanoparticles (GalNP) (Agostini *et al.* 2012). Combination of GalNP with different cargoes offers flexibility to image or eliminate senescent cells (Munoz-Espin *et al.* 2018). However, this flexibility comes to the expense of having to use a modular system, comprised of both the GalNP and the cargo. Here, we propose the use of galactose-modified prodrugs in which a single molecule (the prodrug) is sufficient to target senescent cells taking advantage of their elevated β-galactosidase activity. While we show that duocarmycin derivatives behave as senolytic agents, this approach could be generalised to galactose-modified prodrugs derived of other cytotoxic agents.

In summary, we have described that galactose-modified duocarmycin prodrugs are a new class of broad-spectrum senolytic agents. We have characterized their ability to eliminate different types of senescent cells in culture and *in vivo*. Given the increasing list of diseases that are associated with senescence, galactose-modified duocarmycin prodrugs have the potential to be used in the context of anti-cancer therapies and to treat different age-related diseases.

## EXPERIMENTAL PROCEDURES

### Drugs

The following compounds were used in this study: ABT-263 (Selleckchem, S1001), Etoposide (Sigma-Aldrich, E1383), Q-VD-OPh hydrate (Sigma-Aldrich, SML0063), 4-Hydroxytamoxifen (Sigma-Aldrich, H7904), Doxorubicin hydrochloride (Cayman chemical, 15007). Galactose-modified prodrugs (JHB75B, JHB35B, JHB76B, JHB77B) and seco-Duocarmycin analog dimer were provided by Prof. Dr. L. F. Tietze.

### Antibodies

The following primary antibodies were used in this study: mouse monoclonal anti-BrdU (3D4; BD Biosciences, 555627), mouse monoclonal anti-p16^INK4a^ (JC-8; from CRUK), rabbit polyclonal anti-β-Catenin (Thermo, RB-9035-P1), mouse polyclonal anti-p21 (BD Biosciences, 556431), mouse monoclonal anti-Synaptophysin (27G12; Leica, NCL-L-SYNAP-299), rabbit monoclonal anti-cleaved caspase 3 (Asp175; 5A1E; Cell Signalling Technology, 9664). We used the following secondary antibodies: goat anti-mouse IgG (H+L, AlexaFluor 488 conjugated, Thermo Fischer Scientific, A11029), goat anti-mouse IgG (H+L), AlexaFluor 594 conjugated, Thermo Fischer Scientific, A11032), goat anti-rabbit IgG (H+L, AlexaFluor 594 conjugated, Thermo Fischer Scientific, A11037) and goat anti-rabbit IgG-HRP (Santa Cruz, sc-2004).

### Cell lines

IMR90 cells were obtained from ATCC. IMR90 ER:RAS and IMR90 ER:RAS cells expressing E6 and E7 proteins of HPV16 were generated by retroviral infection of IMR90 cells and have been described elsewhere (Banito *et al.* 2009; Barradas *et al.* 2009). IMR90 were cultured in DMEM (Gibco) supplemented with 10% fetal bovine serum (Sigma) and 1% antibiotic-antimycotic solution (Gibco). HMEC were cultured in Medium 171 (Gibco) supplemented with MEGS (Gibco), 10% fetal bovine serum (Sigma) and 1% antibiotic-antimycotic solution (Gibco). To induce OIS, IMR90 ER:RAS and HMEC ER:RAS were treated with 100 nM 4-hydroxytamoxifen (4OHT, Sigma) reconstituted in DMSO. To induce chemotherapy-induced senescence, IMR90 cells were treated with 0.5 μM Doxorubicin (Sigma) for 24 hours, or with 50 μM Etoposide (Sigma) for 48 hours. To induce senescence by ionizing radiation, IMR90 were γ-irradiated (**6** Gy) and analysed at the indicated times.

### Vector construction

pGIPZ-based shRNA targeting *GLB1* (V3LHS_361778, V3LHS_361779, V2LHS_232389) were obtained from MRC LMS Genomics core facility. To generate IMR90 ER:RAS expressing shRNAs against *GLB1*, lentiviral infections were carried out as described before (Aarts *et al.* 2017). Briefly, HEK293T cells were transfected with the lentiviral and packaging vectors using PEI (PEI 2500, Polysciences). Two days after transfection, HEK293T viral supernatants were collected, filtered (0.45 μM), diluted 1/4, supplemented with 4μg/ml polybrene and added to IMR90 ER:RAS cells plated the day before at a density of 1 million cells per 10 cm dish. Four hours later, lentivirus-containing media was replaced with fresh media. Three days after infection, cells were passaged and cultured for three days in the presence of 1μg/ml puromycin (InvivoGen) to select for infected cells.

### BrdU incorporation

BrdU incorporation assays were performed as previously described (Georgilis *et al.* 2018). Briefly, for BrdU incorporation assays, the cells were incubated with 10 μM BrdU for 16-18 hours before being fixed with 4% PFA (w/v). BrdU incorporation was assessed by Immunofluorescence and High Content Analysis microscopy.

### Immunofluorescence staining of cells

Cells were grown in 96-well plates, fixed with 4% PFA (w/v) and stained as previously described (Georgilis *et al.* 2018).

### Cytochemical SA-β-Galactosidase assay

Cells were grown on 6-well plates, fixed with 0.5% glutaraldehyde (w/v) (Sigma) in PBS for 10-15 min, washed with 1mM MgCl_2_/PBS (pH 6.0) and then incubated with X-Gal staining solution (1 mg/ml X-Gal, Thermo Scientific, 5 mM K_3_[Fe(CN)_6_] and 5 mM K_4_[Fe(CN)_6_] for 8 hr at 37°C. Bright field images of cells were taken using the DP20 digital camera attached to the Olympus CKX41 inverted light microscope. The percentage of SA-β-Gal positive cells was estimated by counting at least 100 cells per replicate sample facilitated by the “point picker” tool of ImageJ software (NIH). For SA-β-Galactosidase staining in tissues, frozen sections (6 μm) were fixed in ice-cold 0.5% glutaraldehyde (w/v) solution for 15 min, washed with 1mM MgCl_2_/PBS (pH 6.0) for 5 min and then incubated with X-Gal staining solution for 16-18 hr at 37°C as previously described (Georgilis *et al.* 2018). After the staining, the slides were counterstained with eosin, dehydrated, mounted and analysed by phase-contrast microscopy. SA-β-Gal tissue staining was quantified using ImageJ software (NIH) by measuring the percentage of stained area in each section and multiplying it by its mean intensity value as described before (Tordella *et al.* 2016). To exclude the luminal spaces in the lung sections, the percentage of SA-β-Gal positive area was divided by the total lung area, as determined by eosin-positive area using ImageJ (NIH).

### Determining senolytic activity

For oncogene-induced senescence experiments, IMR90 ER:RAS or HMEC ER:RAS cells were plated in 96-well dishes and induced to undergo senescence by treating them with 100 nM 4OHT for 6 days. At that point, 1 µM ABT-263 or different concentrations of the galactose-modified pro-drugs were added. In parallel, the same treatments were carried out in IMR90 ER:RAS or HMEC ER:RAS cells treated with DMSO (-4OHT). These cells do not undergo senescence. Cells were fixed at day 9 after 4OHT induction and stained with DAPI (1 μg/ml) for 15 min to assess cell numbers using automated microscopy. Different models of senescence were used to test the senolytic activity of galactose-modified pro-drugs in cell culture in a similar fashion. Briefly, for therapy-induced senescence IMR90 cells were treated with 50 μM etoposide (48h), 0.5 μM doxorubicin (24h) or left untreated, and then kept in drug-free complete media until day 7, when the senolytics were added. Cells were fixed at day 10 after senescence induction. In all senescence types tested, 3-day course of senolytics was applied. The percentage of cell survival was calculated dividing the number of cells after drug treatment by the number of cells treated with vehicle.

### High Content Analysis (HCA)

IF imaging was carried out using the automated high-throughput fluorescent microscope IN Cell Analyzer 2000 (GE Healthcare) with a 20x objective. Multiple fields within a well were acquired in order to include a minimum of 1,000 cells per sample-well. HCA of the images were processed using the INCell Investigator 2.7.3 software as described previously (Herranz *et al.* 2015). Briefly, DAPI served as a nuclear mask hence allowed for segmentation of cells with a Top-Hat method. To detect cytoplasmic staining in cultured cells, a collar of 7-9 μm around DAPI was applied. In samples of cultured cells, a threshold for positive cells was assigned above the average intensity of unstained or negative control sample unless otherwise specified.

### IncuCyte analysis

IMR90 ER:RAS cells were plated in 96-well dishes and induced to undergo senescence as previously described. Different concentrations of galactose-modified pro-drug were added as normally. Cell culture media was supplemented with IncuCyte NucLight Rapid Red reagent for cell labelling (Essen Bioscience) and IncuCyte Caspase-3/7 reagent for apoptosis (Essen Bioscience). Four images per well were collected every 2 hr for 3 days using a 10x objective.

### Mouse models and drug treatments

For induction of senescence, C57BL/6J mice at of 8–12 weeks of age were exposed to a sublethal dose (6 Gy) of total body irradiation. 8 weeks after, mice were injected with 50nmols of pro-drug (i.v.) or vehicle for 4 consecutive days. Mice were killed 24 h after the last injection. Mice lungs were harvested for RNA extraction, paraffin embedded for immunohistology, or frozen in OCT/Sucrose 15% (1:1) solution for cryosectioning and SA-β-gal stains. The mice used for all experiments were randomly assigned to control or treatment groups. Both sexes were used throughout the study.

For in vivo treatment, ABT-263 was prepared in ethanol:polyethylene glycol 400:Phosal 50 PG at 10:30:60 as previously described (Chang *et al.* 2016). Mice were gavaged with vehicle (ethanol:polyethylene glycol 400:Phosal 50 PG) or ABT-263 (50mg/kg). Peripheral whole blood and liver samples were collected 6 hours after dosing.

All mouse procedures were performed under licence, following UK Home Office Animals (Scientific Procedures) Act 1986 and local institutional guidelines (UCL or Imperial College ethical review committees).

### *Ex vivo* culture of mouse pituitaries

Neoplastic pituitaries from 18.5dpc *Hesx1*^*Cre/+*^;*Ctnnb1*^*lox(ex3)/+*^ mice were dissected and placed on top of 0.2 μM Whatman filters (SLS) in 24 well plates containing 500 μl of media (DMEM-F12, Gibco, 1% Pen/Strep, Sigma and 1% FBS, PAA) supplemented with either prodrug A or vehicle (DMSO). Media was changed every 24h, pituitaries were processed for analysis after 72 hours. Immunofluorescence staining was performed as previously described (Gonzalez-Meljem *et al.* 2017). The proportion of β-catenin-accumulating and p21-positive cells was calculated as an index out of the total DAPI-stained nuclei. The proportion of β-catenin-accumulating, cleaved-caspase-3 and p21-positive cells was calculated as an index out of the total DAPI-stained nuclei. Over 300,000 DAPI nuclei were counted from ten histological sections per sample, in a total of twelve neoplastic pituitaries.

### Statistical analysis

GraphPad Prism 7.0 was used for statistical analysis. Two-tailed Student’s *t*-tests were used to estimate statistically significant differences between two groups. Two-way ANOVA with Tukey’s *post hoc* comparison was used for multiple comparisons. Values are presented as mean ± s.d. unless otherwise indicated. Asterisks (*) always indicate significant differences as follows: ns = not significant, * = *P* < 0.5, ** = *P* < 0.01, *** = *P* < 0.001.

For *in vivo* studies, mice were randomly assigned to treatment groups. All replicates in this study represent different mice.

## ACKNOWLEDGEMENTS

We are grateful to members of J. Gil’s laboratory for reagents, comments and other contributions to this project. Core support from MRC (MC-A652-5PZ00 and MC_U120085810) funded this research in J. Gil’s laboratory. D.J.W. was funded by a Wellcome Trust Strategic Award (098565) and core support from MRC (MC-A654-5QB40). J.P.M.-B. was funded by the Brain Tumour Charity (SIGNAL and EVEREST), Children with Cancer UK, CRUK, Great Ormond Street Hospital (GOSH) Children’s Charity and National Institute of Health Research Biomedical Research Centre at GOSH for Children NHS Foundation Trust and University College London. J.P.M.-B. is a GOSH for Children’s Charity Principal Investigator.

## AUTHOR CONTRIBUTIONS

A.G., designed performed and analyzed the cell culture experiments and wrote the first draft of the manuscript. R.G. designed performed and analyzed the experiments with the ACP model. N.H. designed, performed and analyzed the in vivo experiments. L.F.T. designed and synthetized the duocarmycin derivatives and secured funding. A.U., J.P. M.-B. and D.J.W. designed the in vivo experiments and secured funding. J.G conceived and designed the project, secured funding and wrote the manuscript, with all authors providing feedback.

## SUPPORTING INFORMATION LISTING

Supplementary information includes 3 supplementary figures and their legends.

