## Supplementary material for "Galactose-modified duocarmycin prodrugs as senolytics": Sup.

Supplemental information includes 3 supplementary figures and their legends.

Guerrero et al. **Sup. Fig. S1**

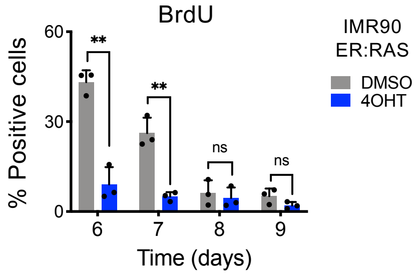


**Supplementary Figure S1. Senescent cells are arrested at the time of treatment with duocarmycin and duocarmycin derivatives.** Quantification of immunofluorescence staining for BrdU. IMR90 ER:RAS were treated with 4-OHT or vehicle (DMSO) for 6 days to induce senescence. Cell were then fixed (day 6) or kept under serum-starvation (0.5% FBS) conditions and fixed 24h (day 7), 48h (day 8) or 72h (day 9) later to assess BrdU incorporation. A 16 h pulse of BrdU was given before fixation ($n$ = 3). Statistical significance was calculated using unpaired two-tailed Student's $t$-tests. Error bars represent mean ± s.d; $n$ represents independent experiments.; ns, not significant; *$P$ < 0.05; **$P$ < 0.01; ***$P$ < 0.001.

Guerrero et al. **Sup. Fig. S2**

a

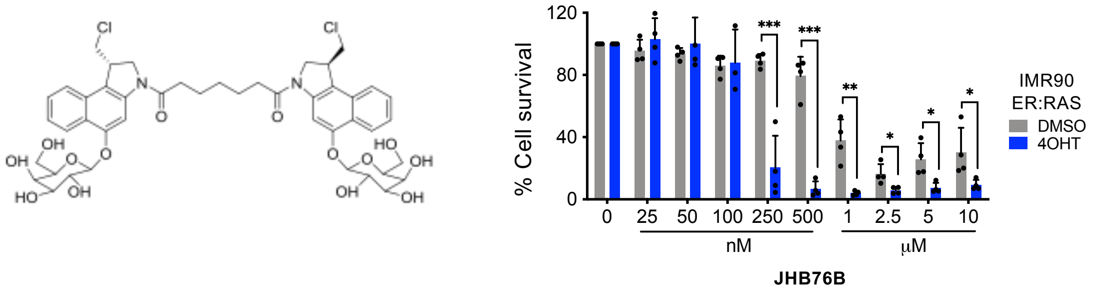


b

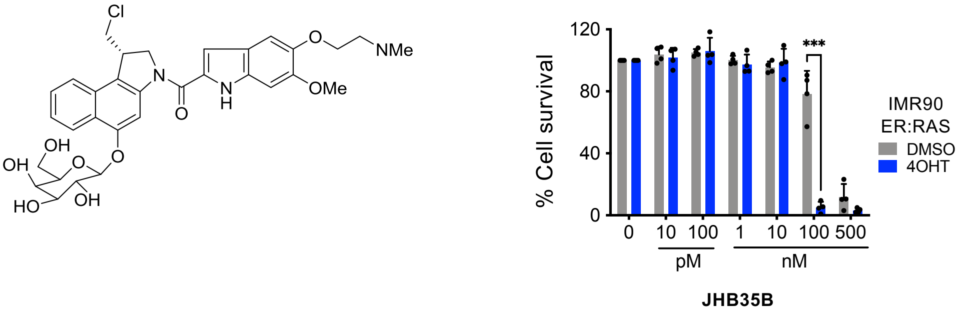


**Supplementary Figure S2. Galactose-modified duocarmycin derivatives selectively kill senescent cells.** Quantification of cell survival after treatment with the galactose-modified duocarmycin derivatives, JHB76B **(a)** and JHB35B **(b)** in the context of oncogene-induced senescence in IMR90 ER:RAS (*n* = 4). A reduced version of this graph is shown in Figure 2f. All statistical significances were calculated using unpaired two-tailed Student's *t*-tests. All error bars represent mean ± s.d; *n* represents independent experiments.; ns, not significant; **P* < 0.05; ***P* < 0.01; ****P* < 0.001.

Guerrero et al. **Sup. Fig. S3**

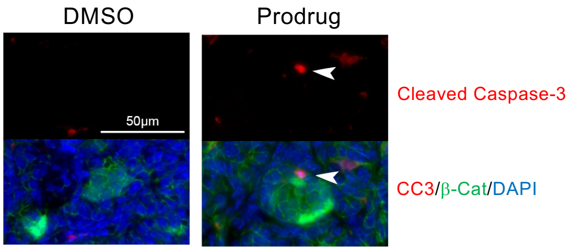


**Supplementary Figure S3. Galactose-modified duocarmycin derivatives induce apoptosis of β-catenin-accumulating preneoplastic cells in a model of ACP.** Representative pictures of pituitaries treated with DMSO (left) or prodrug A (right). Top panels show cleaved caspase-3 staining (red). Bottom panels show a merged composite image showing cleaved caspase-3 staining (red), β-Catenin (green) and DAPI (blue). Scale bar, 50 μM.
